# The conserved upstream ORF of the Arabidopsis *ANAC082* gene mediates translational upregulation in response to nucleolar stress

**DOI:** 10.1101/2022.10.07.511279

**Authors:** Shun Sasaki, Toru Murakami, Miharu Yasumuro, Ayaka Makita, Yutaro Oi, Yuta Hiragori, Shun Watanabe, Rin Kudo, Noriya Hayashi, Iwai Ohbayashi, Munetaka Sugiyama, Yui Yamashita, Satoshi Naito, Hitoshi Onouchi

## Abstract

Perturbations in ribosome biogenesis cause a type of cellular stress called nucleolar or ribosomal stress, which triggers adaptive responses in both animal and plant cells. The Arabidopsis ANAC082 transcription factor has been identified as a key mediator of the plant nucleolar stress response. The 5′-untranslated region (5′-UTR) of *ANAC082* mRNA contains an upstream ORF (uORF) encoding an evolutionarily conserved amino acid sequence. Here, we report that this uORF mediates the upregulation of *ANAC082* translation in response to nucleolar stress. When transgenic Arabidopsis plants containing a luciferase reporter gene under the control of the *ANAC082* promoter and 5′-UTR were treated with reagents that induced nucleolar stress, translation of the reporter gene was enhanced in a uORF sequence-dependent manner. Additionally, we examined the effect of an endoplasmic reticulum (ER) stress-inducing reagent on reporter gene expression because the closest homolog of *ANAC082* in Arabidopsis, *ANAC103*, is involved in the ER stress response. However, the *ANAC082* uORF did not respond to ER stress. Interestingly, although *ANAC103* has a uORF with an amino acid sequence similar to that of the *ANAC082* uORF, the C-terminal sequence critical for regulation is not well conserved among *ANAC103* homologs in Brassicaceae. Transient expression assays revealed that unlike the *ANAC082* uORF, the *ANAC103* uORF does not exert a sequence-dependent regulatory effect. Altogether, our findings suggest that the *ANAC082* uORF is important for the nucleolar stress response but not for the ER stress response, and that for this reason, the uORF sequence-dependent translational regulation was lost in *ANAC103* during evolution.

## Introduction

In eukaryotes, ribosome biogenesis occurs in the nucleolus, which is the largest subnuclear compartment. Perturbations in ribosome biogenesis cause a particular type of cellular stress called nucleolar or ribosomal stress, which is typically accompanied by changes in nucleolar size and morphology (Boulon et al 2010). In animals, nucleolar stress results in cell cycle arrest or apoptosis. As a mechanism of this nucleolar stress response, the pathway involving the tumor suppressor p53 and Murine Double Minute 2 (MDM2), an E3 ubiquitin ligase, has been established (Boulon et al 2010; James et al 2014; Yang et al 2018). Under non-stress conditions, p53 is polyubiquitinated by MDM2 and degraded by the 26S proteasome. When ribosome biogenesis is disturbed and nucleolar stress occurs, unassembled ribosomal proteins are released from the nucleolus into the nucleoplasm. Some of these ribosomal proteins directly bind to MDM2, thereby preventing p53 degradation. Stabilized p53 activates the transcription of many target genes involved in cell cycle arrest and apoptosis, such as the gene encoding p21, an inhibitor of cyclin-dependent kinases (Boulon et al 2010; James et al 2014; Yang et al 2018).

In Arabidopsis, ANAC082, a member of the NAM/ATAF/CUC (NAC) family of transcriptional activators, plays a pivotal role in responses to nucleolar stress (Ohbayashi et al. 2017). Many Arabidopsis mutants deficient in ribosome biogenesis factors (RBFs) or ribosomal proteins have been reported to exhibit altered growth and developmental phenotypes, including embryonic lethality, retarded growth, and abnormal leaf and root development (Horiguchi et al. 2012; Ohbayashi and Sugiyama 2018; Sáez-Vásquez and Delseny 2019; Martinez-Seidei et al. 2020). *ANAC082* was originally identified as the causative gene of the *suppressor of root initiation defective two1 (sriw1)* mutation, which suppressed the cell proliferation defect of the *root initiation defective2 (rid2)* mutant callus (Ohbayashi et al. 2017). The *rid2* mutant is impaired in rRNA processing because of a missense mutation in the gene coding for a putative RNA methyltransferase (Ohbayashi et al. 2011). Additionally, *sriw1/anac082* mutations rescue other growth and developmental phenotypes of ribosome biogenesis-defective mutants, such as retarded growth, narrow and pointed leaves, and abnormalities in root epidermis cell fate (Ohbayashi et al. 2017; Wang et al. 2020). In contrast, the *sriw1* mutation did not rescue the pre-rRNA processing defects in RBF mutants (Ohbayashi et al. 2017). ANAC082 expression is upregulated in response to nucleolar stress (Ohbayashi et al. 2017). Given these observations, ANAC082 is considered to be a key mediator of the nucleolar stress-responsive control of plant growth and development.

We previously identified an upstream ORF (uORF) encoding an evolutionarily conserved amino acid sequence, referred to as a conserved peptide uORF (CPuORF), in the 5′-UTR of *ANAC082* mRNA (Takahashi et al. 2012). This CPuORF has a repressive effect on ANAC082 expression in a peptide sequence-dependent manner (Ebina et al. 2015). Although the sequence conservation across a wide range of angiosperms implies that the *ANAC082* CPuORF has an important regulatory role, its physiological role remains unknown.

In this study, we investigated the possibility that the *ANAC082* CPuORF is involved in nucleolar stress-responsive regulation of ANAC082 expression. Since the closest homolog of *ANAC082* in Arabidopsis, *ANAC103*, is involved in the response to endoplasmic reticulum (ER) stress, we also examined the possibility that the *ANAC082* CPuORF is involved in ER stress-responsive regulation. Our results suggest that the *ANAC082* CPuORF is important for translational regulation in response to nucleolar stress but not to ER stress. Furthermore, we found that although *ANAC103* has a uORF with an amino acid sequence similar to that of the *ANAC082* CPuORF, the *ANAC103* uORF lacks a sequence-dependent regulatory function. We discuss the relationship between the conservation of the *ANAC082* and *ANAC103* uORF sequences and their physiological roles.

## Materials and methods

### Plant materials

*Arabidopsis thaliana* ecotype Col-0 was used for transformation. *A. thaliana* MM2d suspension cells (Menges and Murray 2002) were used for transient expression assays. Cells were cultured in modified Linsmaier and Skoog (LS) medium (Nagata et al. 1992) at 26°C in the dark with orbital shaking at 130 rpm and transferred to fresh medium every week.

### Chemicals

Actinomycin D (ActD) and 5-fluorourasil (5-FU) were purchased from Sigma-Aldrich. Tunicamycin (Tm) was purchased from Funakoshi Co., Ltd.

### Plasmid construction

Plasmid construction is described in Supplementary Text S1. Primers used for plasmid construction are listed in Supplementary Table S1.

### Plant transformation

The *ANAC082pro-5′UTR(WT)::Eluc* reporter construct contains the coding sequence of a Brazilian click beetle luciferase (Emerald luciferase: Eluc) with a PEST degradation sequence (*Eluc-PEST*) under the control of the *ANAC082* promoter and 5′-UTR. Mutant versions of the reporter construct, *ANAC082pro-5′UTR(fs)::Eluc* and *ANAC082pro-5′UTR(Δ ΔΔCPuORF)::Eluc*, contain the frameshift (fs)-mutant CPuORF and the start codon-lacking CPuORF, respectively. Binary vector plasmids carrying each of these reporter constructs were electroporated into *Agrobacterium tumefaciens* strain C58C1Rif^R^ (pGV2260). Arabidopsis plants were transformed with the reporter constructs using the floral dip method (Clough and Bent, 1998). T1 seeds were sown on solid Murashige and Skoog medium supplemented with 50 mg/l kanamycin (Km), 1% (w/v) sucrose, and 0.15% (w/v) agar, and Km-resistant transformants were selected. In the T2 generation, transgenic lines with T-DNA insertions at a single locus or two loci were selected based on the segregation ratios of Km-resistant to Km-sensitive seedlings. Among the transgenic lines obtained, a wild-type reporter line, *ANAC082pro-5′UTR(WT)::Eluc* #1, and the two mutant reporter lines, *ANAC082pro-5′UTR(fs)::Eluc* and *ANAC082pro-5′UTR(Δ ΔΔCPuORF)::Eluc*, were found to have a T-DNA insertion at a single locus. For these, homozygous lines were established based on segregation of Km resistance. Another wild-type reporter line, *ANAC082pro-5′UTR(WT)::Eluc* #2, was found to carry T-DNA insertions at two loci. To establish a homozygous line for both T-DNA insertions by PCR-based genotyping, we determined the T-DNA insertion sites using thermal asymmetric interlaced (TAIL)-PCR (Liu et al., 1995). The homozygous line of *ANAC082pro-5′UTR(WT)::Eluc* #2 was established by genotyping with primers designed based on the T-DNA flanking sequences (Supplementary Table S2).

### Luciferase assay and mRNA quantification in transgenic plants

Approximately 30 seeds of the transgenic plants were surface-sterilized, washed, and grown for six days at 25 °C under continuous light, with shaking at 80 rpm in a 100-ml conical flask containing 10 ml of liquid MGRL medium (Fujiwara et al. 1992). Then, 2.5 or 5 μl of 4 mM ActD, 10 μl of 100 mM 5-FU, or 10 μl of 5 mg/ml Tm was added into the medium. The same amount of dimethyl sulfoxide was added as control treatments. After further incubation for 24 h. the seedlings were frozen in liquid nitrogen and ground with mortar and pestle into a fine powder.

For luciferase assays, approximately 100 mg of the powder was extracted with 200 μl extraction buffer [100 mM (NaH_2_/Na_2_H)PO_4_ and 5 mM dithiothreitol, pH 7] by vortexing. After centrifugation, Eluc activity and protein concentration in supernatant were measured using the Luciferase Assay System (Promega) and the Pierce 660 nm Protein Assay Reagent (Thermo Fisher Scientific), respectively.

For mRNA quantification, total RNA was extracted from approximately 50 mg of the powder using TRIzol (Thermo Fisher Scientific). To remove DNA, 500 ng of total RNAs were treated with 1 unit of RQ1 RNase-Free DNase (Promega) for 30 min at 37 °C. DNase was inactivated by adding 1 μl of RQ1 DNase stop buffer (Promega). The DNase-treated RNAs were subsequently reverse transcribed using SuperScript IV Reverse Transcriptase (Thermo Fisher Scientific) and an oligo (dT) primer, and the synthesized cDNAs were used as templates for quantitative real-time PCR (qRT-PCR). qRT-PCR was performed on a LightCycler 480 System II (Roche Applied Science) using the LightCycler FastStart DNA Master PLUS SYBR Green I kit (Roche Applied Science) following the manufacturer’s instructions. The mRNA levels of *Eluc, ANAC103, UBQ5*, and *ACT2* were measured using the primers listed in Supplementary Table S3.

### Transient expression assay

Transient expression analysis was performed as described by Ebina et al. (2015). Plasmid DNAs were introduced into protoplasts of MM2d suspension cells by electroporation.

## Results

### The *ANAC082* CPuORF mediates nucleolar stress-responsive translational regulation

The amino acid sequence of the *ANAC082* CPuORF is conserved in both dicots and monocots (Figure 1A; Takahashi et al. 2020). The Arabidopsis *ANAC082* CPuORF is 114 nucleotides in length, including the stop codon, and contains two in-frame ATG codons at the first and the 20th codons (Figure 1B). To determine whether the Arabidopsis *ANAC082* CPuORF is involved in the nucleolar stress-responsive regulation of ANAC082 expression, we used transgenic Arabidopsis plants harboring the *Eluc-PEST* reporter gene under the control of the *ANAC082* promoter and 5′-UTR (*ANAC082pro-5′UTR::Eluc*). In addition to the transgenic plants carrying the reporter construct with the wild-type *ANAC082* 5′-UTR [*ANAC082pro-5′UTR(WT)::Eluc*], we generated plants harboring the reporter constructs in which the *ANAC082* CPuORF was mutated. In one of the transgenic reporter plants containing the mutant *ANAC082* CPuORF, the amino acid sequence of the CPuORF in the reporter construct was altered by a fs mutation. This transgenic plant was designated as *ANAC082pro-5′UTR(fs)::Eluc*. In the fs-mutant CPuORF, one nucleotide was inserted between the 68th and 69th nucleotides of the CPuORF, and the 108th nucleotide was deleted (Figure 1B). This fs mutation changed the amino acid residues from the 24th to the 36th position of the CPuORF-encoded peptide but did not alter the CPuORF size (Figure 1B). Our previous study showed that this fs mutation almost completely abolished the repressive effect of the CPuORF on ANAC082 expression (Ebina et al. 2015). In another transgenic reporter plant, *ANAC082pro-5′UTR(ΔCPuORF)::Eluc*, the two ATG codons of the CPuORF were changed to AAG codons to abolish translation of the CPuORF. Using these transgenic plants homozygous for the reporter gene, we first tested the effect of ActD, which is known to induce nucleolar stress by inhibiting the transcription of pre-rRNA (Chen and Hagen 1971; Fraser 1975; James et al 2014), on reporter gene expression. Seeds of the transgenic plants were sown in liquid medium and grown for six days. The seedlings were further incubated for 24 h in the presence or absence of ActD. In experiments using two independent *ANAC082pro-5′UTR(WT)::Eluc* lines, ActD treatment significantly increased reporter gene expression (Figure 2B and C). Measurement of the *Eluc* mRNA level by qRT-PCR showed that the effect of ActD treatment on the mRNA level of the reporter gene was weaker than that on Eluc reporter activity (Figure 2D). Furthermore, the translation efficiency of the reporter gene was calculated by dividing the Eluc activity of each sample by the corresponding *Eluc* mRNA level. As shown in Figure 2E, ActD treatment of *ANAC082pro-5′UTR(WT)::Eluc* seedlings significantly increased the translation efficiency of the reporter gene. These results suggested that the effect of ActD on reporter gene expression was mainly exerted at the translational level. In contrast, treatment of *ANAC082pro-5′UTR(fs)::Eluc* and *ANAC082pro-5′UTR(ΔCPuORF)::Eluc* seedlings with ActD did not significantly affect reporter gene expression (Figure 2B and C), suggesting that the effect of ActD on reporter gene expression depends on the *ANAC082* CPuORF sequence. Additionally, we tested the effect of 5-FU, which induces nucleolar stress by inhibiting pre-rRNA processing (Key 1966; Sun et al. 2007; Wilkinson and Pitot 1973). Treatment of *ANAC082pro-5′UTR(WT)::Eluc* seedlings with 5-FU significantly increased reporter gene expression, whereas 5-FU treatment of the *ANAC082pro-5′UTR(fs)::Eluc* seedlings did not (Figure 3). Altogether, the results of the experiments using ActD and 5-FU suggest that the *ANAC082* CPuORF mediates the upregulation of main ORF (mORF) translation in response to nucleolar stress.

**Figure 1.**
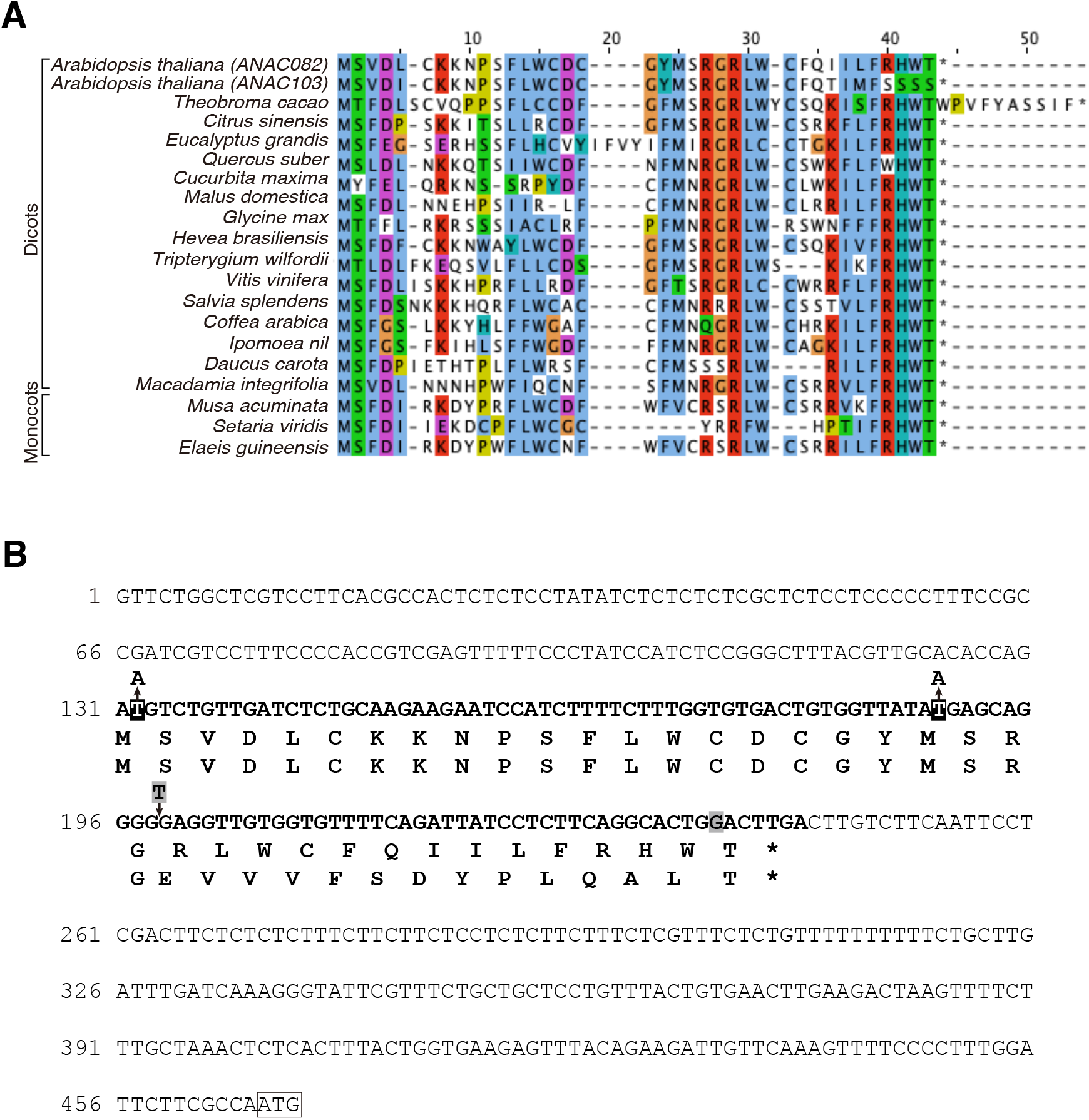
Amino acid sequence conservation of the *ANAC082* CPuORF and the nucleotide sequence of the *ANAC082* 5′-UTR. (A) Alignment of the CPuORF amino acid sequences of *ANAC082* homologs in dicots and monocots. The CPuORF amino acid sequences were aligned using Clustal Omega version 1.2.2 and displayed using Jalview version 2.10.2. The NCBI RefSeq accession numbers of the sequences used to generate the alignment are shown in Supplementary Table S4. (B) Nucleotide sequence of the *ANAC082* 5′-UTR and the deduced amino acid sequence of the CPuORF. The *ANAC082* 5′-UTR nucleotide sequence is based on NCBI RefSeq NM_001085082.1. The nucleotide sequence and the deduced amino sequence of the CPuORF are shown in bold. The nucleotides that were deleted or inserted in the fs mutant are shaded, and the deduced amino sequence of the fs-mutant CPuORF is indicated below that of the wild-type CPuORF. The replaced nucleotides in the *Δ*CPuORF mutant are shown in black background. The AUG initiation codon of the mORF is boxed.

**Figure 2.**
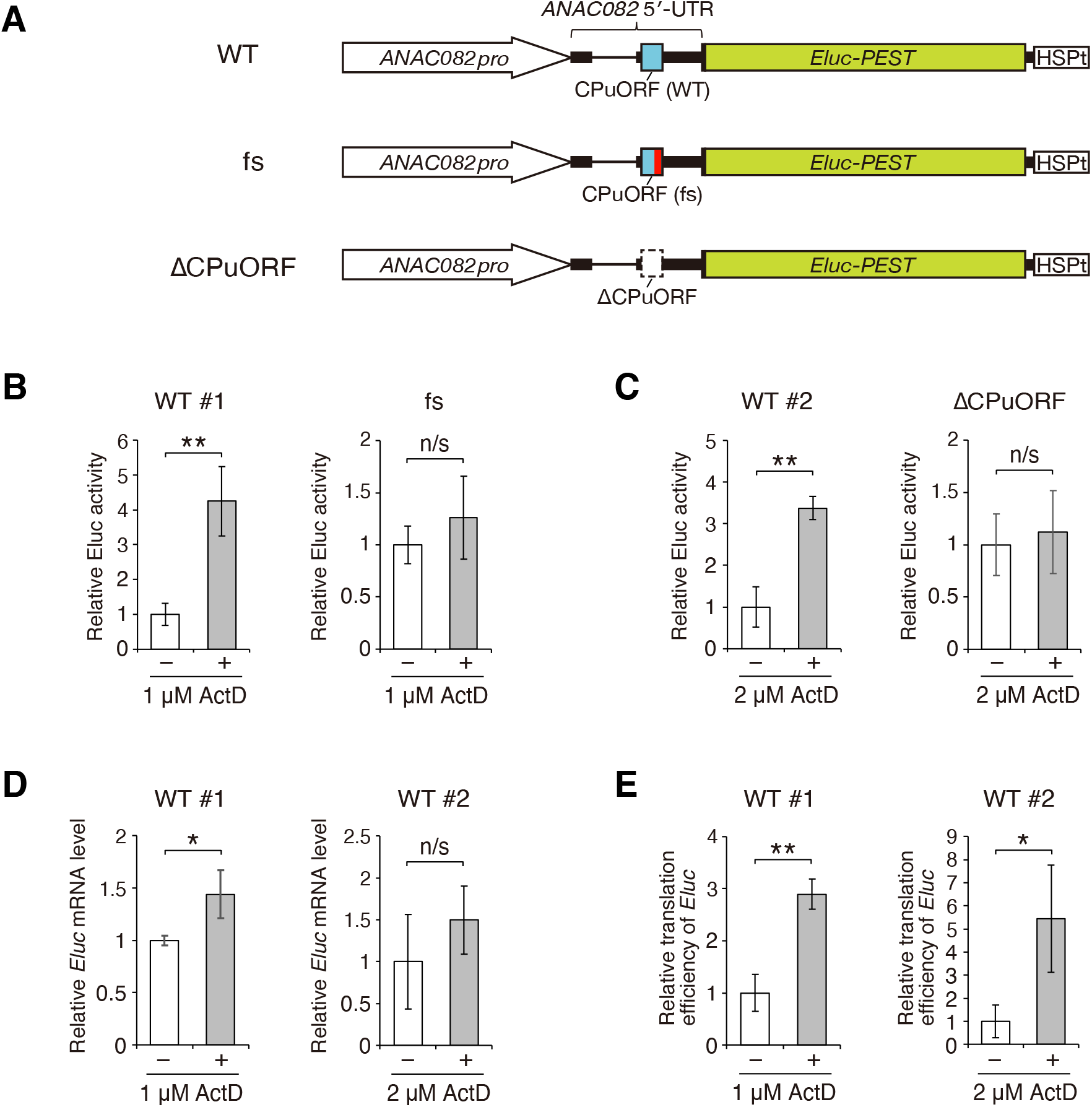
Effect of ActD on CPuORF-mediated regulation of *ANAC082* expression. (A) Schematic representation of the *ANAC082pro-5′UTR::Eluc* constructs. The light blue boxes represent the *ANAC082* CPuORF. The red box in the fs-mutant CPuORF shows the frame-shifted region. The open box with a dotted line depicts the mutant CPuORF lacking the AUG codons (*Δ*CPuORF). The thin line in the *ANAC082* 5′-UTR denotes the intron. *ANAC082pro*, the *ANAC082* promoter; *Eluc-PEST*, the *Eluc-PEST* coding sequence; *HSPt*, the *AtHSP18*.*2* polyadenylation signal. (B and C) Luciferase assays. Seedlings of the *ANAC082pro-5′UTR(WT)::Eluc* lines #1 and #2, an *ANAC082pro-5′UTR(fs)::Eluc* line, and an *ANAC082pro-5′UTR(ΔCPuORF)::Eluc* line were grown in liquid medium for six days and further incubated for 24 h in the absence (−) or presence (+) of 1 or 2 μM ActD, as indicated. The Eluc activity of each sample was normalized to total protein content. (D) Quantification of the mRNA level of the reporter gene. The mRNA levels of *Eluc* and *UBQ5* in the *ANAC082pro-5′UTR(WT)::Eluc* seedlings used in (B) and (C) were measured by qRT-PCR. The *Eluc* mRNA level was normalized to the *UBQ5* mRNA level. (E) Translation efficiency of the reporter gene. The normalized Eluc activity of each sample was divided by the normalized *Eluc* mRNA level in the same sample to calculate the translation efficiency of *Eluc*. In (B) to (E), data represent means ± SD of three biological replicates, relative to those of seedlings not treated with ActD. n/s indicates no significant difference; * and ** indicate significance at *p* < 0.05 and *p* < 0.01, respectively (two-sided Student’s *t*-test).

**Figure 3.**
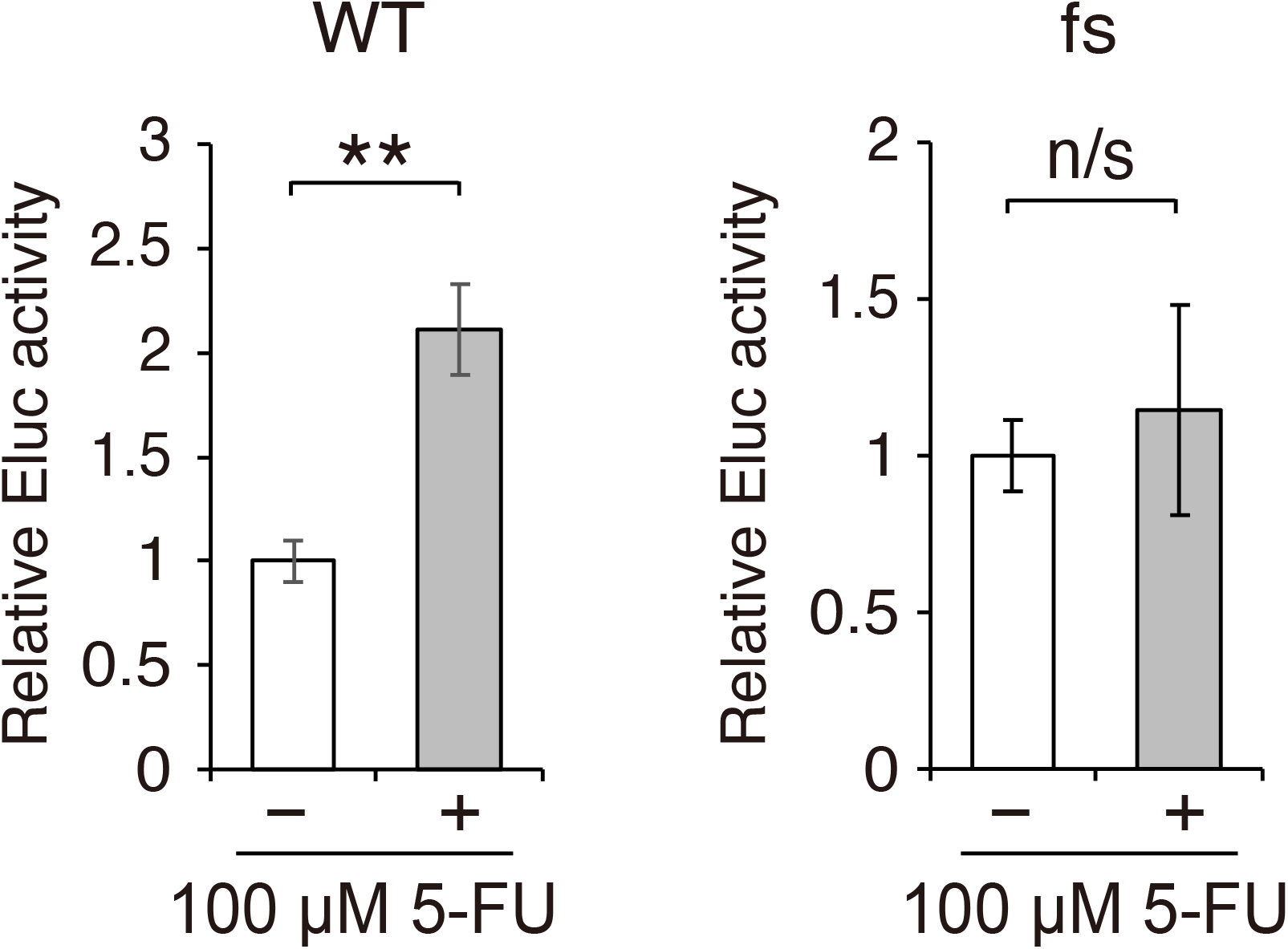
Effect of 5-FU on CPuORF-mediated regulation of *ANAC082* expression. Seedlings of the *ANAC082pro-5′UTR(WT)::Eluc* lines #1 and an *ANAC082pro-5′UTR(fs)::Eluc* line were grown in liquid medium for six days and further incubated for 24 h in the absence (−) or presence (+) of 100 μM 5-FU. The Eluc activity of each sample was normalized to total protein content. Data represent means ± SD of three biological replicates, relative to those of seedlings not treated with 5-FU. n/s indicates no significant difference; ** indicates significance at *p* < 0.01 (two-sided Student’s *t*-test).

### The *ANAC082* CPuORF does not respond to ER stress

ANAC103 is the closest homolog of ANAC082 among the Arabidopsis NAC family of transcription factors, and its 5′-UTR contains a uORF whose amino acid sequence is similar to that of the *ANAC082* CPuORF (Figure 1A). ANAC103 was reported to play a role in the ER stress response (Sun et al. 2013). ER stress is caused by the accumulation of misfolded or unfolded proteins in the ER lumen (Pastor-Cantizano et al. 2020). When ER stress occurs in Arabidopsis plants, *ANAC103* mRNA expression is upregulated (Sun et al. 2013). Therefore, we tested the possibility that the expression of ANAC082 is also upregulated in response to ER stress. To address this possibility, we used Tm, which is known to induce ER stress by inhibiting *N*-linked glycosylation of proteins in the ER (Iwata and Koizumi 2005). As shown in Figure 4A, treatment of *ANAC082pro-5′UTR(WT)::Eluc* and *ANAC082pro-5′UTR(fs)::Eluc* seedlings with Tm did not significantly affect reporter gene expression. In the Tm-treated *ANAC082pro-5′UTR(WT)::Eluc* seedlings, the endogenous *ANAC103* mRNA level was markedly elevated (Figure 4B), as previously reported (Sun et al. 2013). In contrast, the *Eluc* mRNA level did not significantly change, regardless of Tm treatment (Figure 4B). These results suggest that unlike *ANAC103, ANAC082* expression is not regulated in response to ER stress, and that the *ANAC082* CPuORF does not respond to ER stress.

**Figure 4.**
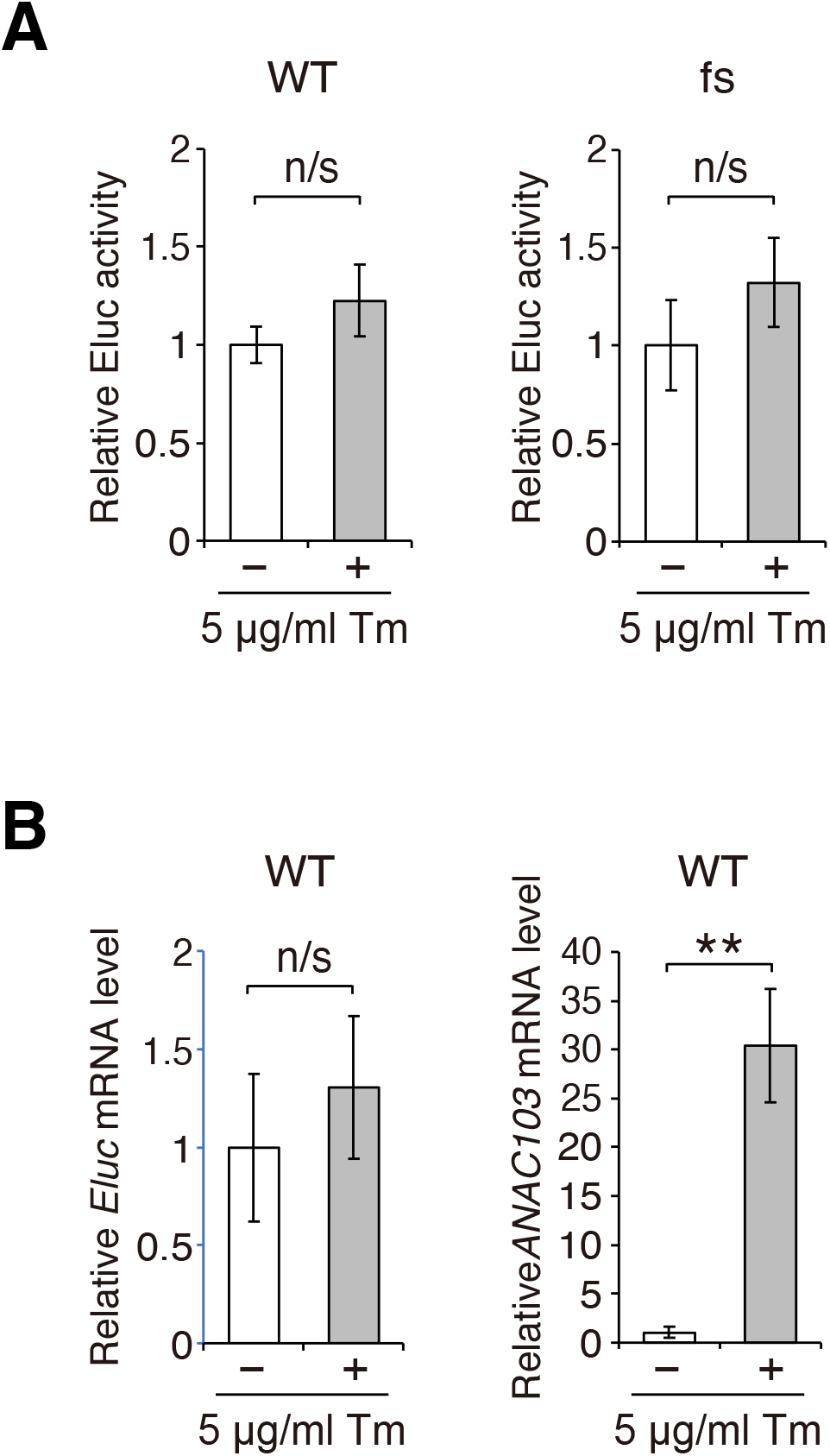
Examination of the effect of Tm on CPuORF-mediated regulation of *ANAC082* expression.(A) Luciferase assays. Seedlings of the *ANAC082pro-5′UTR(WT)::Eluc* lines #1 and an *ANAC082pro-5′UTR(fs)::Eluc* line were grown in liquid medium for six days and further incubated for 12 h in the absence (−) or presence (+) of 5 μg/ml Tm. The Eluc activity of each sample was normalized to total protein content. (B) Quantification of the mRNA levels of the reporter gene and the endogenous *ANAC103* gene. The mRNA levels of *Eluc, ANAC103*, and *ACT2* in the *ANAC082pro-5′UTR(WT)::Eluc* seedlings used in (A) were measured by qRT-PCR. The *Eluc* and *ANAC103* mRNA levels were normalized to the *ACT2* mRNA level. In (A) and (B), data represent means ± SD of at least four biological replicates, relative to those of seedlings not treated with Tm. n/s indicates no significant difference; ** indicates significance at *p* < 0.01 (two-sided Student’s *t*-test).

### The *ANAC103* CPuORF does not exert a sequence-dependent regulatory effect on mORF expression

Although the amino acid sequences of the *ANAC082* and *ANAC103* CPuORFs are homologous, the four C-terminal amino acid residues of the *ANAC103* CPuORF-encoded peptide are different from those of the *ANAC082* CPuORF-encoded peptide (Figure 1A). Considering the lack of response of the *ANAC082* CPuORF to ER stress and the role of ANAC103 in the ER stress response, CPuORF-mediated translational regulation may not be important for ANAC103 expression control and, therefore, may have been lost in *ANAC103* during evolution. To test this hypothesis, we compared the sequence conservation of the *ANAC082* and *ANAC103* CPuORFs in Brassicaceae. As shown in Figure 5A, the CPuORF amino acid sequences are highly conserved among *ANAC082* homologs in Brassicaceae. In contrast, several C-terminal amino acid residues of the CPuORF-encoded peptides are not conserved among *ANAC103* homologs in Brassicaceae (Figure 5A). Our previous study showed that the C-terminal region of the *ANAC082* CPuORF-encoded peptide is critical for CPuORF-mediated translational repression of *ANAC082* (Ebina et al. 2015). Therefore, the low conservation of the CPuORF C-terminal sequences among the *ANAC103* homologs supports the idea that the *ANAC103* CPuORF has lost the regulatory effect in the course of evolution. To further confirm this, we examined whether the *ANAC103* CPuORF lacks the sequence-dependent regulatory effect on mORF expression. For this purpose, we generated the reporter construct in which the *ANAC103* 5′-UTR was fused to the *Renilla* luciferase (Rluc) coding sequence under the control of the cauliflower mosaic virus 35S RNA (35S) promoter (Figure 5C). Then, to determine whether the *ANAC103* CPuORF had a peptide sequence-dependent effect on mORF expression, we mutated the CPuORF in the *ANAC103* 5′-UTR of the reporter construct. In the mutant version of the reporter construct, the amino acid residues from the 23rd to the 28th and from the 34th to the 36th positions of the *ANAC103* CPuORF-encoded peptide sequence were changed by fs mutations (Figure 5C and Supplementary Figure S1). The altered amino acid residues included Arg-24, Trp-26, and Cys-27, which were conserved between the *ANAC082* and *ANAC103* CPuORFs (Figure 1A). In our previous study, when the Arg-24, Trp-26, and Cys-27 codons of the *ANAC082* CPuORF were individually replaced by an alanine codon, all three alanine substitution mutations almost completely abolished the regulatory function of the CPuORF-encoded peptide (Ebina et al. 2015). Therefore, if the *ANAC103* CPuORF-encoded peptide has the conserved regulatory function, the fs mutation in the *ANAC103* CPuORF is expected to abolish the function. For comparison, we also used the reporter constructs in which the wild-type and fs-mutant versions of the *ANAC082* 5′-UTRs were individually fused to the *Rluc* coding sequence under the control of the 35S promoter (Figure 5B) (Ebina et al. 2015). The wild-type and fs-mutant versions of the *ANAC082* and *ANAC103* reporter constructs were separately transfected into protoplasts of *Arabidopsis thaliana* MM2d suspension-cultured cells. After 24 h of incubation, cells were harvested and disrupted to measure luciferase activity. For *ANAC082*, the fs mutation significantly increased reporter gene expression (Figure 5B), as previously observed (Ebina et al. 2015). In contrast, the fs mutation in the *ANAC103* CPuORF did not significantly affect reporter gene expression (Figure 5C). This result suggests that the *ANAC103* CPuORF does not exert a sequence-dependent regulatory effect on mORF expression.

**Figure 5.**
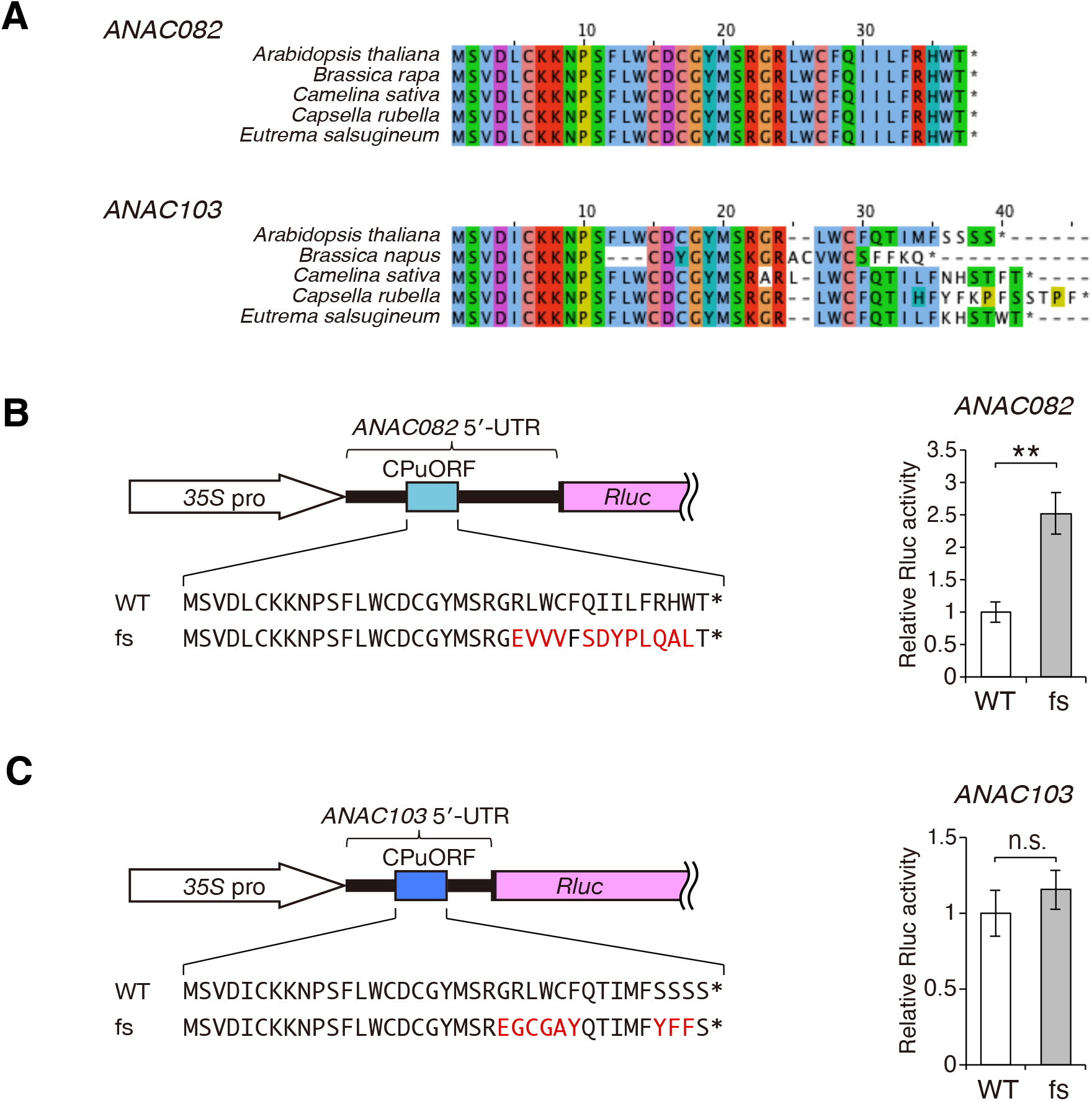
Amino acid sequence conservation of the *ANAC082* and *ANAC103* CPuORFs among homologs in Brassicaceae and examination of the sequence-dependent regulatory function of the *ANAC082* and *ANAC103* CPuORFs. (A) Alignments of the CPuORF amino acid sequences of *ANAC082* and *ANAC103* homologs in Brassicaceae. RefSeq proteins with amino acid sequences most similar to ANAC082 and ANAC103 in a Brassicaceae species were identified by blastp as the closest homologs of ANAC082 and ANAC103 in the species, respectively. Four Refseq RNAs corresponding to the RefSeq proteins were selected for each of *ANAC082* and *ANAC103*, and the CPuORF sequences in the selected Refseq RNAs were used to generate the amino acid sequence alignments of the CPuORFs. The CPuORF amino acid sequences were aligned using Clustal Omega version 1.2.2 and displayed using Jalview version 2.10.2. The RefSeq accession numbers of the sequences used to generate the alignments are shown in Supplementary Table S4. (B and C) Transient expression assays using the reporter constructs containing the *ANAC082* or *ANAC103* CPuORFs. Schematic representation of the reporter constructs is shown at left. The light and dark blue boxes represent the *ANAC082* or *ANAC103* CPuORFs, respectively. The amino acid sequences of the wild-type (WT) and fs-mutant *ANAC082* and *ANAC103* CPuORFs are indicated. The altered sequences in the fs-mutant CPuORFs are shown in red. *35S* pro, the *35S* promoter; *Rluc*, the *Rlu*c coding sequence. Each reporter plasmid containing the WT or fs-mutant versions of *ANAC082* or *ANAC103* CPuORFs was co-transfected into MM2d protoplasts with the *35S::Fluc* internal control plasmid, which carried the firefly luciferase (Fluc) coding sequence under the control of the 35S promoter (Matsuo et al. 2001). The Rluc activity of each sample was normalized to the Fluc activity. Data represent the mean ± SD of the normalized Rluc activities of five biological replicates, relative to that of the corresponding WT reporter construct. n/s indicates no significant difference; ** indicates significance at *p* < 0.01 (two-sided Student’s *t*-test).

## Discussion

ANAC082 was previously reported to play a crucial role in the nucleolar stress response in plants (Ohbayashi et al. 2017). However, the mechanism of nucleolar stress-responsive regulation of ANAC082 expression is unknown. Our study revealed that the ANAC082 expression is translationally upregulated in response to nucleolar stress, and that the CPuORF is responsible for this regulation.

We previously showed that under non-stress conditions, the *ANAC082* CPuORF-encoded peptide represses mORF expression by acting in cis, and that the C-terminal region of the peptide is critical for repression (Ebina et al. 2015). These results suggested that during the translation of *ANAC082* mRNA, the nascent peptide encoded by the CPuORF acts inside the ribosome to repress ANAC082 expression. The most likely mechanism of the CPuORF nascent peptide-mediated repression of ANAC082 expression is that the nascent peptide causes ribosome stalling and the stalled ribosome blocks the access of other scanning ribosomes to the *ANAC082* mORF start codon. This is because several regulatory nascent peptides have been reported to cause ribosome stalling by interacting with components of the ribosomal exit tunnel (Ito and Chiba 2013). The data reported herein revealed that the *ANAC082* CPuORF mediates nucleolar stress-responsive translational regulation in a sequence-dependent manner. This suggests that nucleolar stress alleviates CPuORF peptide-mediated repression of *ANAC082* translation.

Known mechanisms of CPuORF-mediated translational upregulation in response to stress involves phosphorylation of the α subunit of eukaryotic initiation factor 2 (eIF2α). For example, in mammalian *CHOP* mRNA, the translation initiation efficiency of the CPuORF is reduced under ER stress conditions in which eIF2α is phosphorylated. Since the *CHOP* CPuORF-encoded peptide represses translation of the *CHOP* mORF by causing ribosome stalling, the reduced translation efficiency of the CPuORF in response to ER stress leads to ribosomal bypass of the CPuORF and enhanced mORF translation (Palam et al. 2011). Although phosphorylation of eIF2α is induced by various stresses (Baird and Wek 2012; Lokdarshi and von Arnim 2022), nucleolar stress is not known to induce eIF2α phosphorylation in any organism. Given this and our observation that the *ANAC082* CPuORF did not respond to ER stress, it may be unlikely that eIF2α phosphorylation is involved in the CPuORF-mediated translational regulation of *ANAC082*. Further studies are required to determine whether the CPuORF-mediated translational regulation of *ANAC082* in response to nucleolar stress involves eIF2α phosphorylation or some other unknown mechanism.

In animals, upon nucleolar stress, polyubiquitination of p53 by MDM2 is inhibited and p53 is stabilized (Boulon et al 2010; James et al 2014; Yang et al 2018). Interestingly, although the mechanisms of regulation differ, the protein levels of both p53 and ANAC082 are post-transcriptionally upregulated in response to nucleolar stress. This may be because post-transcriptional regulation has an advantage over transcriptional regulation that it enables a more rapid induction of protein expression in response to nucleolar stress. When nucleolar stress occurs, a rapid increase in the expression of these key transcriptional factors may be required to cope with defects in ribosome biogenesis as soon as possible.

Human *MDM2* mRNA contains two uORFs in its 5′-UTR (Jin et al. 2003), and the 5′-proximal uORF is involved in the translational regulation of the *MDM2* mORF in response to hyperosmotic stress (Akulich et al. 2019). However, unlike the *ANAC082* CPuORF, the *MDM2* uORF mediates stress-responsive downregulation of the mORF translation (Akulich et al. 2019). Therefore, the mechanism underlying the uORF-mediated translational regulation differs between *ANAC082* and *MDM2*.

This study showed that unlike the *ANAC082* CPuORF, the *ANAC103* CPuORF does not exert a sequence-dependent effect on mORF expression (Figure 5C). While the C-terminal region of the *ANAC082* CPuORF-encoded peptide is well conserved (Figure 1) and is important for translational repression of the mORF (Ebina et al. 2015), several C-terminal amino acid residues of the CPuORF-encoded peptides of the *ANAC103* homologs are not conserved in Brassicaceae (Figure 5A). Previous studies reported that ANAC103 is involved in the ABA response during seed germination, the ER stress response, and the DNA damage response (Sun et al. 2020; Sun et al. 2013; Ryu et al. 2019). *ANAC103* mRNA expression is up-regulated in response to ABA, ER stress, and DNA damage (Sun et al. 2020; Sun et al. 2013; Ryu 2019). bZIP60, a key transcription factor in the plant ER stress response, directly binds to the *ANAC103* promoter and is required for the ER stress-responsive regulation of *ANAC103* expression (Sun et al. 2013). Similarly, SOG1, a key transcription factor in the plant DNA damage response, also binds directly to the *ANAC103* promoter and is necessary for the DNA damage-responsive *ANAC103* regulation (Ryu et al. 2019). In addition to *ANAC103* mRNA regulation, ER stress and ABA stabilize the ANAC103 protein (Sun et al. 2013; Sun et al. 2020). Thus, *ANAC103* is regulated at the transcriptional and protein stability levels. After an ancestral gene of *ANAC103* diverged from the common ancestor with *ANAC08*2 and acquired stress-responsive transcriptional regulation and protein stability control, stress-responsive translational regulation mediated by the CPuORF may have become less important. Alternatively, since an *ANAC103* ancestor became involved in the ER stress response, the DNA damage response, and the ABA response rather than in the nucleolar stress response, it may no longer have needed translational regulation mediated by the CPuORF that responds to nucleolar stress.

Although this study revealed that the *ANAC082* CPuORF mediates nucleolar stress-responsive translational regulation of the mORF, the mechanisms of how plant cells sense nucleolar stress and how the CPuORF responds to the stress remain to be elucidated. Our findings suggest that the regulatory mechanism for the expression of the key mediator of the nucleolar stress response in plants differs from that in animals. Therefore, plants may have specific mechanisms for sensing and signaling nuclear stress. Further studies are necessary to elucidate these mechanisms.

## Acknowledgements

We thank Ms. Sayo Aono for general assistance. We used the DNA sequencing facility of the Graduate School of Agriculture, Hokkaido University. S.S. and Y.H. acknowledge JSPS for support. Y.Y. is a recipient of the Program for Fostering Researchers for the Next Generation conducted by the Consortium Office for Fostering of Researchers in Future Generations, Hokkaido University.

## Funding

This work was supported by the Japan Society for the Promotion of Science (JSPS) KAKENHI [Grant Nos. JP16H05063 and JP20H05926 to S.N., JP19H02917 and JP19K22299 to H.O., 21K15114 and JP19K16159 to Y.Y,]; the Ministry of Education, Culture, Sports, Science and Technology (MEXT) KAKENHI [Grant Nos. JP17H05658 to S.N.]; the Research Foundation for the Electrotechnology of Chubu to H.T; JSPS Fellows [21J12977 to S.S., 21J21981 to Y.H.]; and the Naito Foundation to H.O.

## Conflict of interest

The authors have no conflicts of interest to declare.

## Description of Supplementary Files

Supplementary Figure S1. Nucleotide sequence of the *ANAC103* 5′-UTRs and the deduced amino acid sequences of the CPuORF

Supplementary Table S1. Primers used for plasmid construction Supplementary

Table S2. Primers used for genotyping

Supplementary Table S3. Primers used for qRT-PCR

Supplementary Table S4. NCBI RefSeq accession numbers of the sequences used to generate the alignments.

Supplementary Text S1. Plasmid construction

## Notes

### Competing Interest Statement

The authors have declared no competing interest.

