## Supplementary data for "The conserved upstream ORF of the Arabidopsis *ANAC082* gene mediates translational upregulation in response to nucleolar stress"

```
  1 CTTCTCCTTCAAGCTATACTCTCATTCTCTCTCCTCCGACCAAAGTTTTCTGTTCCTCCG

 61 CTTCTCCTTCCCCTACCGGTGCTTCGTTTTGTGTTGTACAGCAGATGTCTGTTGATATCT
                                                M  S  V  D  I  C
                                                M  S  V  D  I  C

121 GCAAGAAGAATCCATCGTTTCTTTGGTGTGACTGTGGATATATGAGCAGAGGAAGGTTGT
     K  K  N  P  S  F  L  W  C  D  C  G  Y  M  S  R  G  R  L  W
     K  K  N  P  S  F  L  W  C  D  C  G  Y  M  S  R   E  G  C
           A              A
181 GGTGCTTTCAGACCATCATGTTTTCTTCTTCTTCTTAGCTTTTTGTCTCTGCTAGATCTT
     C  F  Q  T  I  M  F  S  S  S  S  *
    G  A  Y Q  T  I  M  F  Y F  F   S  *

241 GTTATTGTGAGTTTTTTGCTTTACTGGTGAGGGAAGTTACTTAGTGTAGTGAAAGTTTTC

301 CCCTTGTCTTCCAATG
```

**Supplementary Figure S1.** Nucleotide sequence of the ANAC103 5′ -UTR and the deduced amino acid sequence of the CPuORF. The ANAC082 5′ -UTR nucleotide sequence is based on NCBI RefSeq NM_125802.3. The nucleotide sequence and the deduced amino sequence of the CPuORF are shown in bold. The nucleotides that were deleted or inserted in the fs mutant are shaded, and the deduced amino sequence of the fs-mutant CPuORF is indicated below that of the wild-type CPuORF. The AUG initiation codon of the mORF is boxed.

**Supplementary Table S1.** Primers used for plasmid construction

| Primer name | Primer sequence (5′ to 3′) |
|---|---|
| SalI-smGFPf | ATTGTCGACGAGTAAAGGAGAAGAACTTTTCACT |
| smGFP-GUSr | AAGGGACTGACCGGATCCTTTGTATAGTTCATCCATGCCATGT |
| smGFP-GUSf | AAGGATCCGGTCAGTCCCTTATGTTACGTCCTGTAG |
| NLS-SacIR | GTGGAGCTCTTTCCTATCCTCCAACCTTTCTCTTCT |
| XbaI-smGFPf | TCCTCTAGAAAGATGGCGTCGACGAGTAAAGGAGAAGA |
| HSPt-SacIF | CCTGAGCTCGAAGATGAAGATGAAATATTTGGTGTGTCA |
| HSPt-EcoRIR | CCAGAATTCCATAGTCCATACCATAGCACA |
| ElucXBXf | TCTCTCGAGGATCCTCTAGAAAGATGGCGTCGACGGAGAGAGAGAAGAACGTGGTGTAC |
| ElucPESTr | GTGGAGCTCGAATGGCATCTACACATTGATCCT |
| ANAC082proF1 | TCTGGTACCTTTCCATATGAGCGATCGTATGCG |
| ANAC082P2 | TCTGTCGACTTCCCCATTGGCGAAGAATCC |
| ANAC082proF2 | GGCATGCCTCAATATGAAGA |
| ANAC082OLr | GATGGATAGGGAAAAACTCGAC |
| ANAC082OLf | GTCGAGTTTTTCCCTATCCATC |
| ANAC082P3 | TCCGTCGACTTCCCCATTG |
| ANAC103P1 | TCCTCTAGATCTCTCTCCTCCGACCAAAG |
| ANAC103P2 | TCTGTCGACGTTTTCCCCATTGGAAGACAAGG |
| ANAC103fsF | GGTGCTTATCAGACCATCATGTTTTACTTCTTCTCTTG |
| ANAC103fsR | CATGATGGTCTGATAAGCACCACAACCTTCCCTGCTCAT |

**Supplementary Table S2.** Primers used for genotyping

| Primer name | Primer sequence (5′ to 3′) |
|---|---|
| L2GT1Fw | GCATTAACTTGTTGCAGTTGCCTAA |
| L2GT1Rv | GATCCGTTCCGTTCACTACTTGTC |
| L2GT2Fw | AATGGGCCTAGAGCATCTACA |
| L2GT2Rv | CTTGCACCAAGCAGCATGAA |

**Supplementary Table S3.** Primers used for qRT-PCR

| Gene | Forward primer sequence (5′ to 3′) | Reverse primer sequence (5′ to 3′) |
|---|---|---|
| *Eluc* | GAGAGCCTGCACAACTTCA | GTTCCTGTGGGTCTGCAT |
| *ANAC103* | CCACCTCTAACAAGTGATGTTATAGC | CCATCAGGATGTCTTACCTCA |
| *UBQ5* | GTGGTGCTAAGAAGAGGAAGA | TCAAGCTTCAACTCCTTCTTT |
| *ACT2* | CTTGCACCAAGCAGCATGAA | CCGATCCAGACACTGTACTTCCTT |

**Supplementary Table S4.** NCBI RefSeq accession numbers of the sequences used to generate the alignments.

| Figure number | Species | RefSeq accession numbers |
|---|---|---|
| Figure 1A | *Arabidopsis thaliana (ANAC082)* | NM_001085082.1 |
| | *Arabidopsis thaliana (ANAC103)* | NM_125802.3 |
| | *Theobroma cacao* | XM_007048279.2 |
| | *Citrus sinensis* | XM_006464580.3 |
| | *Eucalyptus grandis* | XM_018862738.1 |
| | *Quercus suber* | XM_024028567.1 |
| | *Cucurbita maxima* | XM_023147648.1 |
| | *Malus domestica* | NM_001328926.1 |
| | *Glycine max* | XM_003526813.4 |
| | *Hevea brasiliensis* | XM_021825146.1 |
| | *Tripterygium wilfordii* | XM_038830077.1 |
| | *Vitis vinifera* | XM_003631867.3 |
| | *Salvia splendens* | XM_042165189.1 |
| | *Coffea arabica* | XM_027222650.1 |
| | *Ipomoea nil* | XM_019321701.1 |
| | *Daucus carota* | XM_017373971.1 |
| | *Macadamia integrifolia* | XM_042639279.1 |
| | *Musa acuminata* | XM_009421903.2 |
| | *Setaria viridis* | XM_034727913.1 |
| | *Elaeis guineensis* | XM_029262078.1 |
| Figure 5A (*ANAC082*) | *Arabidopsis thaliana* | NM_001085082.1 |
| | *Brassica rapa* | XM_009124259.3 |
| | *Camelina sativa* | XM_010424710.2 |
| | *Capsella rubella* | XM_006287843.2 |
| | *Eutrema salsugineum* | XM_006399344.2 |
| Figure 5A (*ANAC103*) | *Arabidopsis thaliana* | NM_125802.3 |
| | *Brassica napus* | XM_013787594.3 |
| | *Camelina sativa* | XM_010463058.2 |
| | *Capsella rubella* | XM_006280618.2 |
| | *Eutrema salsugineum* | XM_006394145.2 |

### Supplementary Text S1

#### Materials and methods

#### Plasmid construction

Plasmid pBI-A082WT-Eluc carries the *ANAC082pro-5′UTR(WT)::Eluc* reporter construct, which contains the *Eluc-PEST* coding sequence under the control of the *ANAC082* promoter and 5′-UTR, in the binary vector pBI121 (Clontech Laboratories). This plasmid was constructed as follows. The coding sequence of the soluble modified red shifted green fluorescent protein was amplified by PCR from pBI-Ex1-GFP (Suzuki et al. 2001) with primers SalI-smGFPf and smGFP-GUSr. On the other hand, the β-glucuronidase (GUS) coding sequence with the nuclear localization signal (NLS) of the SV40 large T-antigen was amplified from the 35S-sGFP-GUS-NLS construct (Matsushita et al. 2003) with primers smGFP-GUSf and NLS-SacIR. These two PCR fragments were fused by overlap extension PCR (Ho et al. 1989) with primers XbaI-smGFPf and NLS-SacIR. The fused fragment was digested with XbaI and SphI and ligated into XbaI/SphI-digested pGEM-7Zf+ (Promega) to yield plasmid pTME1. pTME1 was digested with HindIII and SacI, and the resulting 2,640-base-pair fragment was ligated into HindIII/SacI-digested pBI121 to generate plasmid pTME2. Then, the region containing the polyadenylation signal of the *HSP18.2* gene was amplified from *A. thaliana* (Col-0 ecotype) genomic DNA by PCR with primers HSPt-SacIF and HSPt-EcoRIR. The amplified fragment was digested with SacI and EcoRI and ligated into SacI/EcoRI–digested pTME2 to make plasmid pTME3. The *Eluc-PEST* coding sequence was amplified from pELuc(PEST)-test (Toyobo) with primers ElucXBXf and ElucPESTr. The amplified fragment was digested with KpnI and SacI and ligated into KpnI/SacI–digested pTME3 to yield plasmid pKXBX-Eluc. The region containing the *ANAC082* promoter and 5′-UTR was amplified from *A. thaliana* Col-0 genomic DNA by PCR with primers ANAC082proF1 and ANAC082P2. The amplified fragment was digested with KpnI and SalI and ligated into KpnI/SalI–digested pKXBX-Eluc to make plasmid pBI-A082WT-Eluc.

Plasmids pBI-A082fs-Eluc and pBI-A082ΔCPuORF-Eluc carry mutant versions of the

*ANAC082pro-5′UTR::Eluc* reporter constructs in pBI121 and contain the start codon-lacking CPuORF and the fs-mutant CPuORF, respectively. To generate pBI-A082fs-Eluc, the region containing the *ANAC082* promoter and a 5′ part of the *ANAC082* 5′-UTR was amplified from pBI-A082WT-Eluc with primers ANAC082proF2 and ANAC082OLr. On the other hand, the *ANAC082* 5′-UTR containing the fs mutant CPuORF was amplified from the fs-mutant version of the *35S::ANAC082-UTR:RLUC* reporter plasmid (Ebina et al. 2015) with primers ANAC082OLf and ANAC082P3. These two PCR fragments were fused by overlap extension PCR with primers ANAC082proF2 and ANAC082P3. The fused fragment was digested with SpeI and SalI and ligated into SpeI/SalI-digested pBI-A082WT-Eluc. To create pBI-A082ΔCPuORF-Eluc, the *ANAC082* 5′-UTR was amplified from the ΔAUG-mutant version of the *35S::ANAC082-UTR:RLUC* reporter plasmid (Ebina et al. 2015) with primers ANAC082OLf and ANAC082P2. This PCR fragment was fused 3′ to the above-mentioned fragment containing the *ANAC082* promoter and a 5′ part of the *ANAC082* 5′-UTR, using overlap extension PCR with primers ANAC082proF2 and ANAC082P2. The fused fragment was digested with SpeI and SalI and ligated into SpeI/SalI-digested pBI-A082WT-Eluc.

Plasmid pEIANAC103 harbors the cauliflower mosaic virus 35S RNA (35S) promoter, the *ANAC103* 5′-UTR, the *Renilla* luciferase (Rluc) coding sequence, and the polyadenylation signal of the *Agrobacterium tumefaciens* nopaline synthase (*nos*) gene in pUC19. To construct this plasmid, we amplified the *ANAC103* 5′-UTR by reverse transcription PCR from total RNA of seven-day-old *A. thaliana* Col-0 seedlings, using the OneStep RT-PCR Kit (Qiagen) with primers ANAC103P1 and ANAC103P2. The amplified fragment was digested with XbaI and SalI and ligated into the XbaI/SalI-digested pIE0 (Ebina et al. 2015). Mutations were introduced into the *ANAC103* CPuORF of pEIANAC103 using overlap extension PCR with primers ANAC103fsF and ANAC103fsR.

Primers used for plasmid construction are listed in Supplementary Table S1. Sequence analysis confirmed the integrity of the PCR-amplified regions of all constructs.
